# Offline RL for generative design of protein binders

**DOI:** 10.1101/2023.11.29.569328

**Authors:** Denis Tarasov, Ulrich A. Mbou Sob, Miguel Arbesú, Nima Siboni, Sebastien Boyer, Marcin Skwark, Andries Smit, Oliver Bent, Arnu Pretorius

## Abstract

Offline Reinforcement Learning (RL) offers a compelling avenue for solving RL problems without the need for interactions with an environment, which may be expensive or unsafe. While online RL methods have found success in various domains, such as *de novo* Structure-Based Drug Discovery (SBDD), they struggle when it comes to optimizing essential properties derived from protein-ligand docking. The high computational cost associated with the docking process makes it impractical for online RL, which typically requires hundreds of thousands of interactions during learning. In this study, we propose the application of offline RL to address the bottleneck posed by the docking process, leveraging RL’s capability to optimize non-differentiable properties. Our preliminary investigation focuses on using offline RL to conditionally generate drugs with improved docking and chemical properties.

## 1 Introduction

The process of designing new drugs is extremely challenging, given the required investment of time and resources required for a drug to navigate through the various stages, spanning from its conception and development, pharmacological characterization and rigorous testing, and eventual approval for mass production and clinical use [15]. Furthermore, the vast chemical space comprising all potential drug candidates and their intricate interactions with other molecules and proteins pose formidable obstacles [27]. During the design phase in SBDD, our goal is to identify drug candidates with a higher likelihood of successfully traversing these stages – i.e. ensuring favorable chemical attributes such as synthetic accessibility or drug-likeness, while also targeting specific disease-related protein targets as selectively as possible.

Recent advancements in artificial intelligence (AI) have provided valuable tools to aid researchers in the quest for better drug candidates [1]. Some of these tools are dedicated to assessing molecular properties[25, 38], enabling the validation of molecules before embarking on costly real-world experiments. Another avenue of research focuses on generating molecules themselves [13, 16, 26]. Many such approaches rely on deep learning techniques, necessitating differentiability for training. However, most molecular properties are rule-based or predicted using diverse models, rendering gradient-based learning infeasible for such signals. This challenge can be effectively addressed by incorporating techniques from Reinforcement Learning (RL).

RL has exhibited promising results across a wide range of domains, including robotics [43] and Natural Language Processing (NLP)[29]. Online RL techniques have also found application in drug generation tasks [26]. However, the necessity for direct interaction with the environment can be a substantial bottleneck for RL algorithms, given the associated costs and potential risks, particularly when algorithms require thousands or even millions of interactions. In SBDD, molecular docking is a critical process that assesses whether a molecule interacts with a target protein, and in this case serves as a prime example of such a costly interaction. Incorporating this property in the *de novo* molecule generation process is key to achieve successful drug candidates [8].

Offline RL methods [22] leverage the power of RL without the need for direct interaction with the environment, relying solely on pre-collected datasets of interactions. Numerous successful applications of offline RL exist in fields like robotics[31, 20], autonomous driving[10], and recommendation systems[7]. In this work, our primary objective is to apply offline RL techniques to the domain of *de novo* drug design, where drugs must satisfy a predefined set of chemical and docking properties.

Drawing inspiration from the successes of RL in the field of NLP, our proposed approach seeks to generate SMILES [40] strings for potential drugs, conditioned on the binding site of the target protein. To achieve this, first we train the generative model on the supervised task of generating sequences similar to the dataset, after that we employ an offline RL methodology, aiming to enhance the quality of generated samples in terms of docking and chemical properties.

This preliminary work serves as a proof of concept, demonstrating the potential of utilizing offline RL to address intricate optimization challenges in drug generation that are difficult to tackle using conventional approaches. Additionally, we propose a strategy for data augmentation, aiming to solve the limited availability of structural protein-ligand complex data. This approach paves the way for future research endeavours in this area.

## 2 Related work

The domain of AI-driven drug generation has witnessed substantial research progress[5]. Previous approaches to generating drugs have leveraged a variety of neural network architectures. These encompass variants of Variational Autoencoders (VAE) [13, 16], Recurrent Neural Networks (RNN) [28], Graph Neural Networks (GNN) [42], Generative Adversarial Networks (GAN) [14], and Transformers [3, 26].

Given the necessity to generate molecules with specific properties, the application of RL for drug design has gained traction in recent years [28, 6, 39, 41, 2, 26]. These RL-based approaches have demonstrated efficacy in optimizing one or multiple chemical properties that are computationally inexpensive to evaluate. However, the optimization of docking properties — which instead rely or expensive physics-based simulation — remains a formidable challenge [8].

It’s noteworthy that some research endeavors have harnessed offline RL techniques in sensitive domains like personalized patient treatment [30, 12]. Intriguingly, as of our knowledge cutoff, there have been no prior attempts to apply offline RL to the conditional *de novo* drug design.

## 3 Preliminaries

### 3.1 Docking Process in Drug Design

The docking process is a pivotal step in SBDD. It plays a crucial role in predicting whether a specific molecule, often referred to as a ligand, can effectively interact with a pharmacological target — in most instances, a protein. This interaction is fundamental to the identification of potential drug candidates and development of novel pharmaceutical compounds.

Docking involves the precise prediction of how a ligand (drug) molecule fits into the binding site or active pocket of a target protein. The objective is to assess whether the ligand can form stable and biologically relevant interactions with the protein binding site, often mimicking the natural binding of endogenous ligands or substrates. To do so, it is required to place 3D representations of the target (obtained experimentally) and the ligand under a force field. The initial ligand pose in the binding site is then optimized under the force field so, an energy-minimized structure the bound complex is obtained. The structural properties — e.g. position of the ligand *vs* the pocket, number of interacting atoms — and the relative energy of the complex can be then used to score its stability.

The docking process is an indispensable tool in computational drug design, enabling researchers to screen vast libraries of molecules efficiently and identify potential drug candidates. However, it is computationally intensive and can be a time-consuming aspect of drug discovery, making it an important target for optimization through advanced techniques such as offline reinforcement learning, as explored in this study.

### 3.2 Language Modeling

We leverage the potent encoder-decoder Transformer architecture [37] as the foundation for our generative model. This architecture comprises stacked encoder and decoder blocks, with the decoder blocks structured similarly to the encoder but incorporating certain inputs from the encoder blocks. Each of these blocks relies on the attention mechanism, a pivotal component in the Transformer framework. The attention mechanism operates on keys (*K*), queries (*Q*), and values (*V* ), and its calculation is defined as follows:

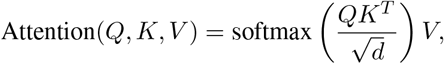

here, *d* represents the dimensionality of the representation. In the encoder, *K* and *V* are derived from the outputs of the preceding blocks, while in the decoder, they are obtained from the final encoder block.

The rationale behind selecting the Transformer architecture for our task is grounded in its remarkable track record of success across diverse domains. Transformers have achieved exceptional performance in various applications, and they have recently shown promise in the field of drug generation [3, 26]. Furthermore, we want to leverage the recently released ESM2 [23] transformer model to compute embeddings for the target proteins on which our generative model is conditioned.

### 3.3 RL Task Formulation

Drawing inspiration from the work of [32], we formalize our problem as a Partially Observable Markov Decision Process [33]. At each timestep *t*, the agent (i.e. the generative model) receives an observation that consist of all previously generated tokens in our sequence i.e. *o*^*t*^ = {*o*^1^, *o*^2^, *…, o*^*t−*1^ }, and predicts the next token for which its receives a reward *r*^*t*^. In our specific context, tokens correspond to the set of all possible atoms and bonds present in our chosen SMILES’s vocabulary. A special *begining of string* token is used at the start to indicate we are generating the first token and another special *end of string* token is used to indicate the end of the sequence. Given that we can only compute the properties of the molecule when the full sequence is transformed to the molecule, all timesteps have reward 0 except the final timestep.

Our formulation can be summarised as follows:

1. Generative model: given some target information *𝒯* and a ligand with SMILES’s representation *{o*^1^, *o*^2^, *…, o*^*n*^*}*, find a model *π*_*θ*_ such that

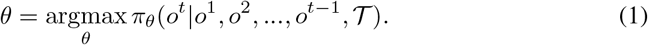
2. RL: finetune the generative model to maximise the expected return ℛ

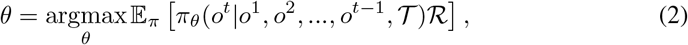

where *ℛ* = *γ*^*t*^*Σ r*^*t*^ and *γ* is the traditi onal discount factor in reinf orcement learning.

## 4 Method

In this section, we outline our method, which comprises three fundamental steps: data preparation, supervised training of the generative model, and offline RL training. These steps collectively form the core of our approach and are pivotal to achieving our research objectives.

### 4.1 Data

#### 4.1.1 Supervised Data

In our research, we harnessed the PDBBind dataset [24], a resource that supplies triplets of 3D structures comprising a protein, a binding pocket, and a ligand. Notably, the inclusion of information about the pocket is a distinctive feature of the PDBBind dataset, rendering it the sole viable option for our SBDD task.

For our SBDD approach, we need string representations of both proteins and ligands. Biopython [9] was used to transform proteins strudtures into amino acids sequences and the binding pockets into binary masks to indicate the presence or not of each amino acids in the binding pocket. For the ligands, we used RDKit [21] to generate SMILES strings, representing their molecular structures. SMILES representation pose a challenge in molecular generation because of the difficultly to accurately capture branches and rings. Additionally, a single molecule can have multiple SMILES representations. Hence, for training we use a recently developed molecular string representation called SELFIES [19] that is more robust than SMILES, in that any random sequence of SELFIES tokens can be synthesised into a valid molecule. We convert all the SMILES strings in our dataset to their SELFIES representations and after training the generated SELFIES are converted back to their equivalent SMILES sequences.

#### 4.1.2 RL Data

Preparing the data for RL finetuning requires scoring the various ligands. For this purpose, we use the following metrics:

1. **Quantitative Estimate of Drug-likeness (QED)**: The QED is a measure of the likelihood of a generated molecule to have molecular properties similar to known drugs [4].
2. **Synthetic Accessibility Score (SAS)**: The SAS is a metric that measures the level of difficulty to synthesize the generated molecule.
3. **Number of satisfied Lipinski’s rules (Lipinski)**: Lipsinki’s rules are a set of guidelines used to assess the likelihood of the generated molecule being suitable for use as a drug based on physichochemical parameters.
4. **Retrosynthetic Accessibility Score (RAS)**: This metric estimates whether it is possible to identify a retrosynthetic path for obtaining the compound [35].

Each score was subsequently mapped to the range [0, 1]. In addition to these property scores, we utilized a GPU-accelerated version [11] of the Vina [36] algorithm to obtain docking scores for the ligands. For each ligand, we first converted it from SELFIES to a 3D molecule using RDKit^2^. Subsequently, we dock the ligand to the target protein using Vina, initializing the position at the center of masses of the corresponding pocket. We recorded the following metrics related to the docking process:

1. **VinaE**: Vina energy score of the best resulting configuration.
2. **Dist**: Distance between the centers of mass of the ligand and the pocket.
3. **Clash**: Number of instances where the ligand atoms and proteins are too close such that a large repulsive force will be generated that will make the complex unstable.

It’s important to note that solely relying on the VinaE score is insufficient to gauge the quality of docking for the target pocket. The distance to the pocket is also crucial in this regard, as it can report wether the docking algorithm could not fit the ligand inside the binding site. Similarly, the number of clashes can be potentially indicative of physically impossible docking configurations.

Following the removal of samples that RDKit or Vina failed to process, the dataset’s final size was 17,133 samples, down from the original 19,447. Running the docking process for each dataset sample took approximately 100 hours in total, utilizing Quadro-RTX-4000 GPUs, averaging to 21 seconds per sample. This extensive computational effort underscores our motivation for optimizing the docking property through offline RL.

#### 4.1.3 Data Augmentation

The dataset size of 17,133 samples is a relatively small for the complex task of drug generation. Unfortunately, the supervised portion of our dataset, which necessitates experimentally determined triplets of proteins, pockets, and ligands specifically binding to the target pockets of proteins, cannot be readily augmented.

However, we can augment the RL datasets by introducing additional docking examples. To achieve this, we randomly select pairs of proteins and pockets and endeavor to dock random ligands from the dataset, following the same process detailed in subsubsection 4.1.2. To prevent redundancy, we exclude triplets that already exist in the PDBBind dataset. This augmentation strategy exposes the model not only to positive examples of ligand binding but also to negative ones, as we assume that a random ligand has a low likelihood of successful docking to a randomly chosen protein. This assumption is supported by the distribution plots shown in Figure 1 and the statistics presented in Table 1.

**Table 1:**
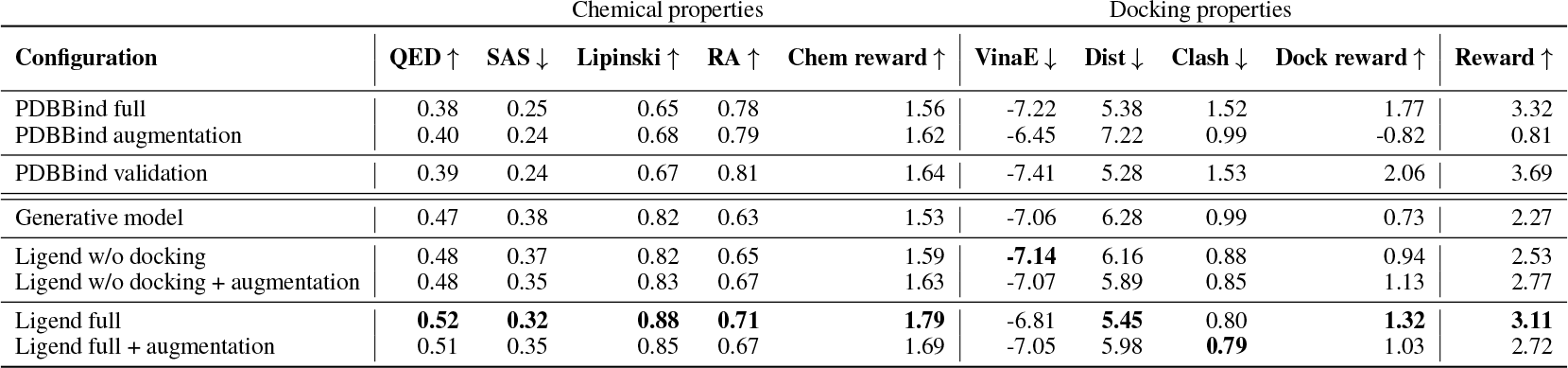
Average results of scoring on the intersection of successfully processed validation sets. *↓* and *↑* indicates that metric must be minimized or maximized respectively. In “w/o docking” all the docking metrics are removed from the reward function. “+ augmentation” indicates the usage of the augmented data. Chemical properties are normalized into the range [0, 1]. We also report the same metrics for the entire PDBBind dataset and it’s augmentation. We highlight the best of scores for generated molecules with the **bold text**.

| Configuration | Chemical properties |  |  |  |  | Docking properties |  |  |  |  |
| --- | --- | --- | --- | --- | --- | --- | --- | --- | --- | --- |
| | QED ↑ | SAS ↓ | Lipinski ↑ | RA ↑ | Chem reward ↑ | VinaE ↓ | Dist ↓ | Clash ↓ | Dock reward ↑ | Reward ↑ |
| PDBBind full | 0.38 | 0.25 | 0.65 | 0.78 | 1.56 | -7.22 | 5.38 | 1.52 | 1.77 | 3.32 |
| PDBBind augmentation | 0.40 | 0.24 | 0.68 | 0.79 | 1.62 | -6.45 | 7.22 | 0.99 | -0.82 | 0.81 |
| PDBBind validation | 0.39 | 0.24 | 0.67 | 0.81 | 1.64 | -7.41 | 5.28 | 1.53 | 2.06 | 3.69 |
| Generative model | 0.47 | 0.38 | 0.82 | 0.63 | 1.53 | -7.06 | 6.28 | 0.99 | 0.73 | 2.27 |
| Ligend w/o docking | 0.48 | 0.37 | 0.82 | 0.65 | 1.59 | **-7.14** | 6.16 | 0.88 | 0.94 | 2.53 |
| Ligend w/o docking + augmentation | 0.48 | 0.35 | 0.83 | 0.67 | 1.63 | -7.07 | 5.89 | 0.85 | 1.13 | 2.77 |
| Ligend full | **0.52** | **0.32** | **0.88** | **0.71** | **1.79** | -6.81 | **5.45** | 0.80 | **1.32** | **3.11** |
| Ligend full + augmentation | 0.51 | 0.35 | 0.85 | 0.67 | 1.69 | -7.05 | 5.98 | **0.79** | 1.03 | 2.72 |

**Figure 1:**
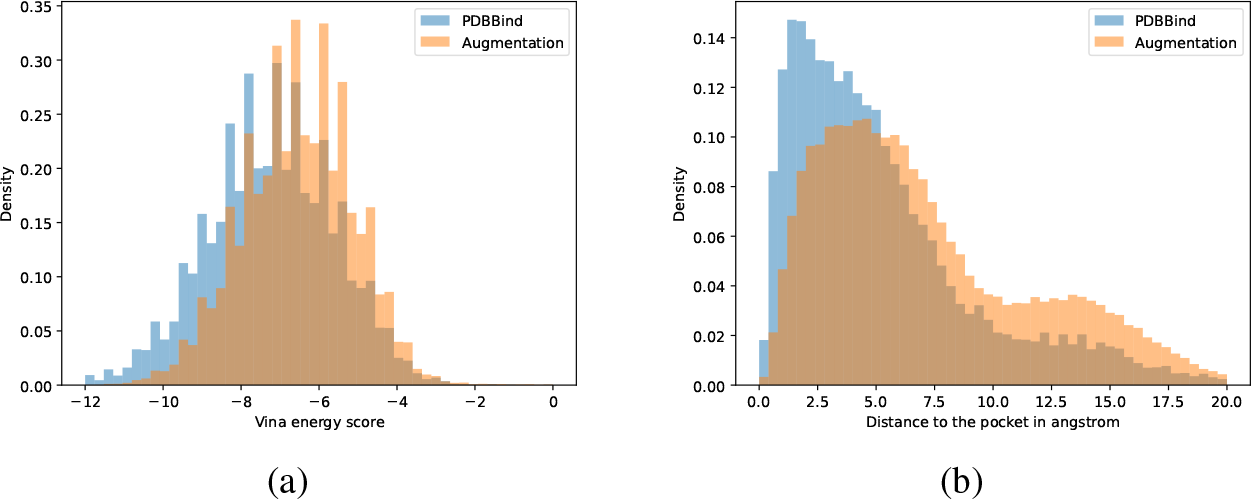
Distribution densities of (a) Vina energy score and (b) distance to the target pocket for PDBBind dataset and its augmentation. The lower values of both metrics correspond to better docking results.

The result of this data augmentation effort is a dataset comprising 77,344 samples. It’s worth noting that this number can be further increased, albeit at the cost of significantly higher computational resources required for docking.

### 4.2 Generative model

The initial training stage of our method focuses on training a generative model using the supervised objective, leveraging the PDBBind dataset. In this stage, we provide the target protein sequence and the binding site mask as inputs to the encoder. The decoder’s task is to predict the next token based on the encoder’s output and all previous tokens in the sequence. We utilize cross-entropy loss as the training objective.

Notably, our approach introduces a novel concept: explicit conditioning on the protein and binding site, a factor not previously employed in attempts to address the binding problem. Our hypothesis is that this conditioning mechanism may enhance the generation of relevant drugs and facilitate better generalization.

To enhance the representation learning for proteins, given the limited dataset size, we opt to utilize the ESM2 protein encoder [23], a pre-trained encoder that has proven effective in capturing protein features. In this regard, we do not retrain this component of our encoder. However, to incorporate binding site information, we introduce additional trainable parameters into our encoder. The decoder is trained from scratch. The supervised training stage of our method is illustrated at the top of Figure 2.

**Figure 2:**
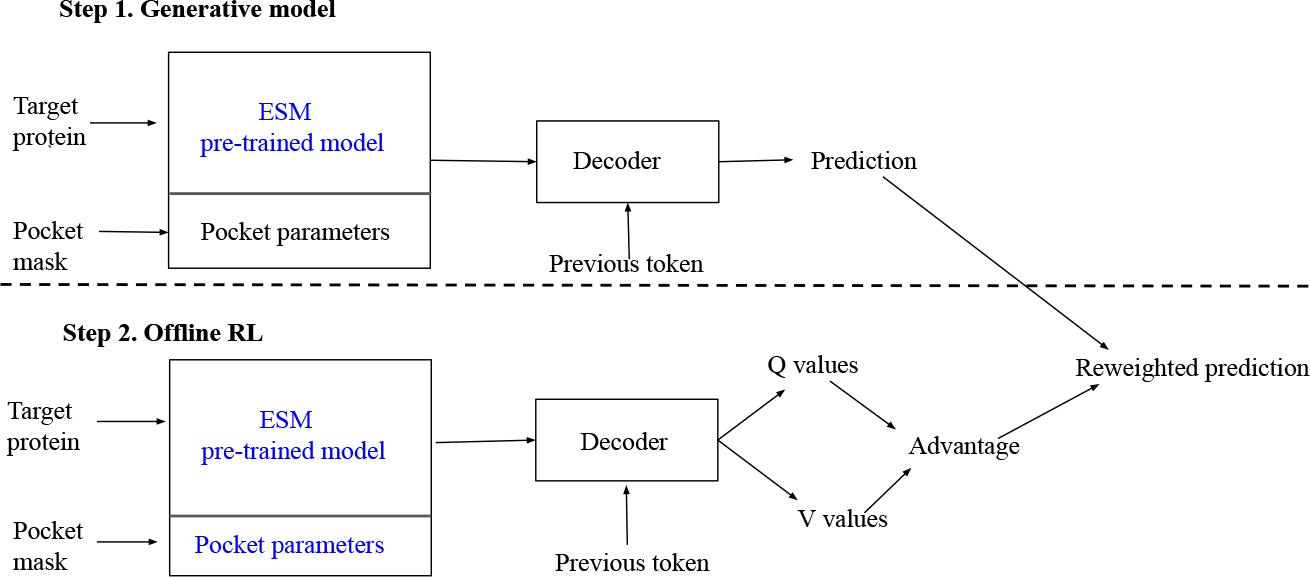
Schema of the proposed approach. Top part of the figure shows the generative model stage. Bottom part of the figure shows the offline RL stage. Text in blue indicates frozen model parameters at various stages.

### 4.3 Offline RL

For the offline RL component of our method, we draw inspiration from ILQL [32], initially proposed for addressing NLP problems. ILQL is built upon the foundation of IQL [18], an offline RL algorithm recognized as one of the most potent offline RL methodologies to date [34]. Additionally the algorithm is very easy to implement.

The underlying concept of ILQL involves taking a pre-trained decoder and replacing the prediction head with a state value function *V* and an action state value function *Q*. In value based RL, the *V* and *Q* functions for a state *s* and an action *a* are defined as follows respectively:

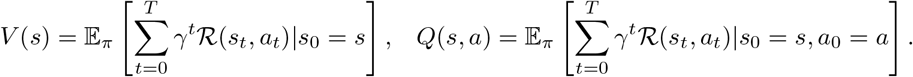

The *V* value provides an estimate of the expected discounted reward that a policy will accumulate when starting from a given state *s*. It offers insights into the overall value of being in a specific state. On the other hand, the *Q* function estimates the expected discounted reward after taking a specific action *a* in a particular state *s*. During training we use the ILQL expectile loss as defined in [32] to learn the *V* and *Q* functions.

During model inference, we employ the outputs of the generative model and shift the predicted token distribution with an advantage obtained from the *V* and *Q* functions, utilizing a weighting parameter *β*. The formula for the resulting score of token *a* given the input *h* (comprising the protein, pocket, and previous tokens) is expressed as:

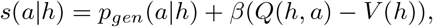

where *p*_*gen*_(*a* |*h*) represents the probability of token *a* predicted by the generative model. The offline RL stage is visually represented at the bottom of Figure 2.

At present, our reward function is a weighted combination of the factors outlined in subsubsection 4.1.2. The weights can be found in subsection A.2.

We christen our approach as **Ligend**, short for Ligand Generation via Structural Coditioning, encapsulating our novel methodology.

## 5 Experiments

### 5.1 Experimental Setup

Using our approach we train models with different configurations in order to see which components are essential and which are not. The following parts of the method were probed: using or not using data augmentation and using or not using docking properties.

We split the PDBBind dataset into training and validation sets. The validation dataset size is only 5% which results into 800 samples due to two factors. First, original dataset size is small, so removing a big part of it may result in severe model performance drop. Second, the docking process required for the evaluation takes a lot of time, so the increase of the validation set size increases the computational overhead.

First, we train the generative model with the training subset of PDBBind. After that, for each of the experiment we reuse the obtained model for further RL tuning and use the same split of the PDBBind for the training.

We evaluate each of the resulting models with the same set of *β* hyperparameters. We choose the best model based on the highest average reward it achieves on validation set. For the list of hyperparameters see subsection A.2.

After collecting predictions for all of the best models we use the intersection of predictions sets which were successfully processed with evaluation pipeline for fair comparison. This results into 554 samples.

### 5.2 Experimental Results

The outcomes of our method under various configurations are presented in Table 1, showcasing a breakdown of each reward component and the cumulative rewards for chemical properties, docking properties, and the overall score.

It is important to note that we did not conduct a hyperparameter search while RL performance may improve strongly with better parameters.

Several noteworthy insights can be gleaned from our results. Firstly, and somewhat surprisingly, the generative model produces predictions for molecules with relatively favorable chemical properties when compared to the dataset distribution. Additionally, the average VinaE score demonstrates notable performance. It must be remarked that the PDBBind full and validation sets are different from the augmented in a fundamental sense: the experimental X-ray structures represent rigid, energy-minimized states not only for the ligand, but also the targets’ binding pocket residues. This means that the structure of the target is optimized towards its bound ligand in the PDBBind dataset. Therefore, it is not surprising that the VinaE scores are significantly better— and that the newly generated molecules can not reach the same energy range — as well as the larger presence of clashes, an artifact from the crystallization. Interestingly, the average VinaE is similar between the Ligend models and the generative one. Optimizing solely for chemical properties yields marginal improvements in chemical scores but comes at the cost of a significant decline in docking results.

When we employ all metrics simultaneously for optimization, a consistent enhancement in chemical properties is observed, alongside a boost in docking properties. Interestingly, the physichochemicalbased properties QED and Lipinski rule of 5 show the smallest improvement over the generative model — suggesting that the learned distribution captures druglikeness — while the SAS and RA, related to the more abstract concept of synthetizability, show the larger changes. Regarding the docking parameters, what distinguishes the full model from the rest, is the reduced average drug-pocket distance and number of clashes, which may compensate the slightly less favorable average dockeing energy.

It is remarkable that the improvements across all metrics were often smaller when the augmentation dataset is included. This behavior with the augmented data might be attributed to the influence of the CQL loss component within the ILQL loss, which forces to elevate the scores for samples from the dataset. In the augmented dataset, we encounter approximately four times as many negative examples of docking compared to positive instances, potentially contributing to this observed effect. Addressing this issue may be possible through more refined hyperparameter selection.

Optimizing for both chemical and docking properties simultaneously leads to improved chemical scores. This improvement reaffirms the idea that molecular docking involves an interplay of various chemical properties which need to be optimised during docking along with the traditional docking specific metrics such as the docking distance or the energy score. Nevertheless, the most significant conclusion remains that RL can effectively enhance predictions compared to those of the generative model improving it by approximately 50%. Despite this improvement, a considerable gap persists between the samples from PDBBind and the output generated by Ligend.

To ascertain the diversity and novelty of the molecules generated by our approach, we provide details in Table 2, encompassing the count of unique molecules, the number of novel molecules not present in the entirety of PDBBind and the Tanimoto similarity between the Morgan fingerprints of generated samples and the molecules in the PDBBind dataset. It is worth noting a slight decrease in the count of unique and novel molecules after RL finetuning compared to the Generative model, indicating a potential avenue for future exploration. Both the Generative and RL finetuned models have average Tanimoto similarities below 0.4, hence both models do not just generate molecules with minor differences compared to those in the datasets but attempt to generate molecules with completely different structures.

**Table 2:** Number of unique molecules, number of molecules which are not present in the PDBBind dataset, and mean Tanimoto similarity between Morgan fingerprints with molecules in the PDBBind dataset, computed using RDKit.

| Configuration | Unique molecules ↑ | Novel molecules ↑ | Fingerprints similarity ↓ |
| --- | --- | --- | --- |
| PDBBind full | 12131 | N/A | N/A |
| PDBBind validation | 446 | N/A | N/A |
| Generative model | 485 | 467 | 0.38 |
| Ligend w/o docking | 479 | 465 | 0.38 |
| Ligend w/o docking + augmentation | 471 | 456 | 0.38 |
| Ligend full | 452 | 443 | 0.36 |
| Ligend full + augmentation | 468 | 456 | 0.36 |

## 6 Conclusion and Future Work

In this study, we have demonstrated a proof of concept, showcasing the potential of offline reinforcement learning (RL) in the domain of *de novo* SBDD, with a focus on specific chemical and docking properties.

A significant limitation of our work is that RL, by nature, optimizes the reward specified, which may not fully capture the complexities of real-world applications. Even the optimal policy derived under a given reward function may produce solutions that are impractical or unsuitable for actual use. To address this limitation, additional expert scoring of the generated molecules is essential to ensure the practical viability of the solutions.

While we have achieved promising results, they remain several avenues for improvement that have the potential to significantly enhance the performance of our approach:

1. **Hyperpararemters** search. Our current experimentation did not involve hyperparameter tuning. For the supervised learning step, this may be less critical, but fine-tuning hyperparameters in the RL phase could substantially boost performance, even in the absence of other enhancements.
2. **Better generative model pretraining**. Given the small dataset size, achieving the challenging generative objective requires additional considerations. An additional pretraining stage, preceding the current supervised phase, could potentially lead to better performance. This new stage could involve generating arbitrary molecules without specific conditioning, utilizing datasets significantly larger than PDBBind. Existing research in this area provides pre-trained models that can be leveraged without the need for extensive training.
3. **Advanced decoding techniques**. In this work, we utilized greedy decoding for generating predictions. However, more sophisticated techniques like beam search are known to offer improved performance. Implementing advanced decoding methods could yield further gains.
4. **Enhanced reward design**. The design of the reward function is a crucial aspect of RL applications. Our current reward function is a simple weighted sum of various metrics, with weights that have not been tuned (except for setting them to zero in ablations). Exploring nonlinear combinations of these metrics may also lead to substantial performance improvements. Also, adding additional scores such as toxicity might be advantageous.

We anticipate that our further work will refine and optimize our approach. We hope that our research will provide valuable insights and inspiration for fellow researchers in the field of drug generation with specific properties.

## Acknowledgments and Disclosure of Funding

Research supported with Cloud TPUs from Google’s TPU Research Cloud (TRC).

## A Appendix

## A.1 Sample generated molecules

Figure 3 shows molecules generated for 20 randomly selected proteins for a visual comparison. Notably, some of the generated molecules feature large rings with > 10 members, often unsaturated. This could be attributed to our model’s limitation in fully capturing the various branching patterns inherent in SMILES molecule representations, leading to the formation of these large rings instead of the more common 5-to 8-member rings. In addition, some of the molecules carry ring fusions subject to unrealistic strains. The model does however make use of common moieties, like phosphate, sulphate, or trifluoromethyl groups, as well as other halogens as Cl.

**Figure 3:**
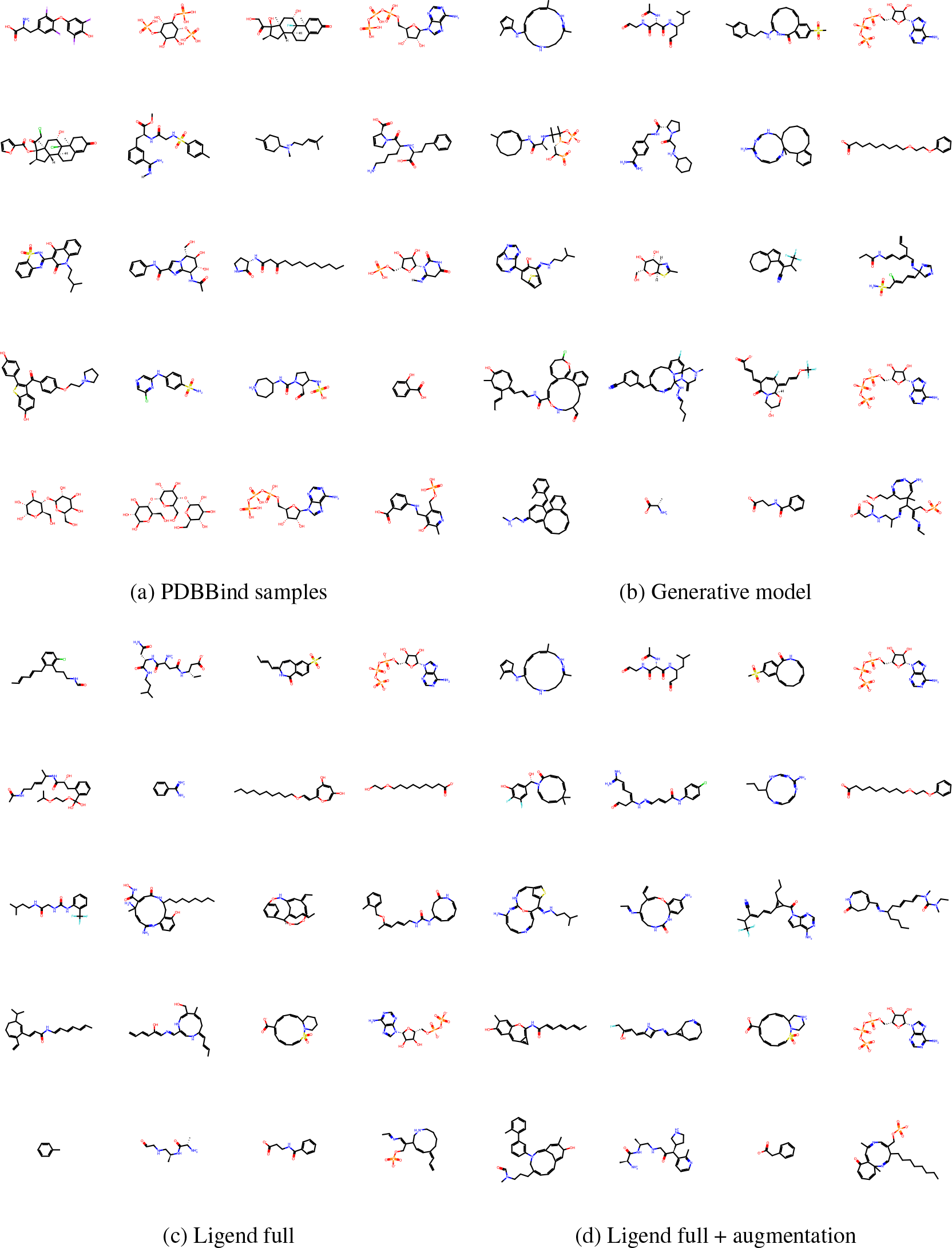
Example molecules generated for 20 randomly selected proteins from the dataset. (a) PDBBind, (b) Generative model, (c) Ligend full and (d) Ligend full + augmentation.

### A.2 Training and inference parameters

We provide the reward parameters and training hyperparameters in Table 3.

**Table 3:** Ligend hyperparameters.

|  | Hyperparameter | Value |
| --- | --- | --- |
| Chemical reward weights | QED | 1.0 |
|  | SAS | -1.0 |
|  | Lipinski | 1.0 |
|  | RA | 1.0 |
| Docking reward weights | VinaE | -1.0 |
|  | Dist | -1.0 |
|  | Clash | -0.05 |
| Generative model | Batch size | 128 |
|  | Optimizer | Adam [17] |
|  | Learning rate | 1e-4 |
|  | Training epochs | 1500 |
| ILQL hyperparameters | Batch size | 128 |
|  | Optimizer | Adam [17] |
|  | Learning rate | 3e-4 |
|  | Training epochs | 100 |
|  | Discount factor | 0.97 |
|  | Target update rate | 5e-3 |
| | IQL $\tau$ | 0.6 |
|  | V loss weight | 1.0 |
|  | Q loss weight | 1.0 |
|  | CQL loss weight | 0.1 |

For inference, we systematically explored multiple *β* parameters, which govern the magnitude of RL influence, from a predefined set of values: 1.0, 3.0, 6.0, 10.0. The optimal *β* values employed to obtain the results presented in Table 1 are detailed in Table 4.

**Table 4:**
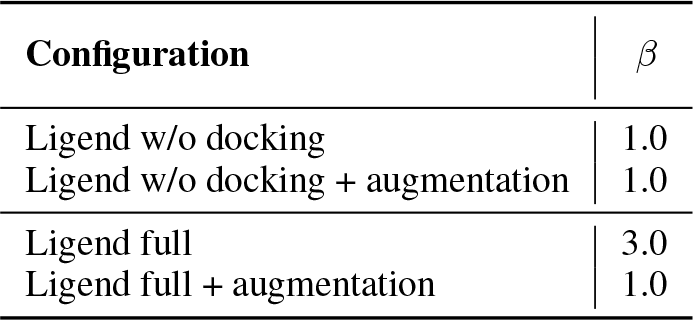
Best *β* values for different Ligend configurations.

| Configuration | $\beta$ |
| --- | --- |
| Legend w/o docking | 1.0 |
| Legend w/o docking + augmentation | 1.0 |
| Legend full | 3.0 |
| Legend full + augmentation | 1.0 |

### Footnotes

2 Note, that we translate the string back to 3D structure and not using the original structure because for newly generated molecules we do not have 3D structures which we want to dock.

## References

[1] Óscar Álvarez-Machancoses and Juan Luis Fernández-Martínez. Using artificial intelligence methods to speed up drug discovery. Expert opinion on drug discovery, 14(8):769–777, 2019.

[2] Sara Romeo Atance, Juan Viguera Diez, Ola Engkvist, Simon Olsson, and Rocío Mercado. De novo drug design using reinforcement learning with graph-based deep generative models. Journal of Chemical Information and Modeling, 62(20):4863–4872, 2022.

[3] Viraj Bagal, Rishal Aggarwal, PK Vinod, and U Deva Priyakumar. Molgpt: molecular generation using a transformer-decoder model. Journal of Chemical Information and Modeling, 62(9): 2064–2076, 2021.

[4] G. R. Bickerton, G. V. Paolini, J. Besnard, S. Muresan, and A. L. Hopkins. Quantifying the chemical beauty of drugs. Nat Chem, 4(2):90–98, Jan 2012.

[5] Camille Bilodeau, Wengong Jin, Tommi Jaakkola, Regina Barzilay, and Klavs F Jensen. Generative models for molecular discovery: Recent advances and challenges. Wiley Interdisciplinary Reviews: Computational Molecular Science, 12(5):e1608, 2022.

[6] Thomas Blaschke, Josep Arús-Pous, Hongming Chen, Christian Margreitter, Christian Tyrchan, Ola Engkvist, Kostas Papadopoulos, and Atanas Patronov. Reinvent 2.0: an ai tool for de novo drug design. Journal of chemical information and modeling, 60(12):5918–5922, 2020.

[7] Minmin Chen, Can Xu, Vince Gatto, Devanshu Jain, Aviral Kumar, and Ed H. Chi. Off-policy actor-critic for recommender systems. Proceedings of the 16th ACM Conference on Recommender Systems, 2022.

[8] Tobiasz Cieplinski, Tomasz Danel, Sabina Podlewska, and Stanislaw Jastrzebski. Generative models should at least be able to design molecules that dock well: A new benchmark. Journal of Chemical Information and Modeling, 2023.

[9] Peter JA Cock, Tiago Antao, Jeffrey T Chang, Brad A Chapman, Cymon J Cox, Andrew Dalke, Iddo Friedberg, Thomas Hamelryck, Frank Kauff, Bartek Wilczynski, et al. Biopython: freely available python tools for computational molecular biology and bioinformatics. Bioinformatics, 25(11):1422–1423, 2009.

[10] Christopher P. Diehl, Timo Sievernich, Martin Krüger, Frank Hoffmann, and Torsten Bertram. Umbrella: Uncertainty-aware model-based offline reinforcement learning leveraging planning. ArXiv, abs/2111.11097, 2021.

[11] Ji Ding, Shidi Tang, Zheming Mei, Lingyue Wang, Qinqin Huang, Haifeng Hu, Ming Ling, and Jiansheng Wu. Vina-gpu 2.0: Further accelerating autodock vina and its derivatives with graphics processing units. Journal of Chemical Information and Modeling, 63(7):1982–1998, 2023.

[12] Harry Emerson, Matthew Guy, and Ryan McConville. Offline reinforcement learning for safer blood glucose control in people with type 1 diabetes. Journal of Biomedical Informatics, 142:104376, 2023.

[13] Rafael Gómez-Bombarelli, Jennifer N Wei, David Duvenaud, José Miguel Hernández-Lobato, Benjamín Sánchez-Lengeling, Dennis Sheberla, Jorge Aguilera-Iparraguirre, Timothy D Hirzel, Ryan P Adams, and Alán Aspuru-Guzik. Automatic chemical design using a data-driven continuous representation of molecules. ACS central science, 4(2):268–276, 2018.

[14] Gabriel Lima Guimaraes, Benjamin Sanchez-Lengeling, Carlos Outeiral, Pedro Luis Cunha Farias, and Alán Aspuru-Guzik. Objective-reinforced generative adversarial networks (organ) for sequence generation models. arXiv preprint arXiv:1705.10843, 2017.

[15] James P Hughes, Stephen Rees, S Barrett Kalindjian, and Karen L Philpott. Principles of early drug discovery. British journal of pharmacology, 162(6):1239–1249, 2011.

[16] Wengong Jin, Regina Barzilay, and Tommi Jaakkola. Junction tree variational autoencoder for molecular graph generation. In International conference on machine learning, pages 2323–2332. PMLR, 2018.

[17] Diederik P Kingma and Jimmy Ba. Adam: A method for stochastic optimization. arXiv preprint arXiv:1412.6980, 2014.

[18] Ilya Kostrikov, Ashvin Nair, and Sergey Levine. Offline reinforcement learning with implicit q-learning. arXiv preprint arXiv:2110.06169, 2021.

[19] Mario Krenn, Florian Häse, AkshatKumar Nigam, Pascal Friederich, and Alan Aspuru-Guzik. Self-referencing embedded strings (selfies): A 100% robust molecular string representation. Machine Learning: Science and Technology, 1(4):045024, 2020.

[20] Aviral Kumar, Anikait Singh, Stephen Tian, Chelsea Finn, and Sergey Levine. A workflow for offline model-free robotic reinforcement learning. In 5th Annual Conference on Robot Learning, 2021. URL https://openreview.net/forum?id=fy4ZBWxYbIo.

[21] Greg Landrum. Rdkit documentation. readthedocs, 1(1-79):4, 2013. URL https://buildmedia.readthedocs.org/media/pdf/rdkit/latest/rdkit.pdf.

[22] Sergey Levine, Aviral Kumar, George Tucker, and Justin Fu. Offline reinforcement learning: Tutorial, review, and perspectives on open problems. arXiv preprint arXiv:2005.01643, 2020.

[23] Zeming Lin, Halil Akin, Roshan Rao, Brian Hie, Zhongkai Zhu, Wenting Lu, Nikita Smetanin, Allan dos Santos Costa, Maryam Fazel-Zarandi, Tom Sercu, Sal Candido, et al. Language models of protein sequences at the scale of evolution enable accurate structure prediction. bioRxiv, 2022.

[24] Zhihai Liu, Minyi Su, Li Han, Jie Liu, Qifan Yang, Yan Li, and Renxiao Wang. Forging the basis for developing protein–ligand interaction scoring functions. Accounts of chemical research, 50 (2):302–309, 2017.

[25] Andreas Mayr, Günter Klambauer, Thomas Unterthiner, and Sepp Hochreiter. Deeptox: toxicity prediction using deep learning. Frontiers in Environmental Science, 3:80, 2016.

[26] Eyal Mazuz, Guy Shtar, Bracha Shapira, and Lior Rokach. Molecule generation using transformers and policy gradient reinforcement learning. Scientific Reports, 13(1):8799, 2023.

[27] Pavel G Polishchuk, Timur I Madzhidov, and Alexandre Varnek. Estimation of the size of drug-like chemical space based on gdb-17 data. Journal of computer-aided molecular design, 27:675–679, 2013.

[28] Mariya Popova, Olexandr Isayev, and Alexander Tropsha. Deep reinforcement learning for de novo drug design. Science advances, 4(7):eaap7885, 2018.

[29] Rajkumar Ramamurthy, Prithviraj Ammanabrolu, Kianté Brantley, Jack Hessel, Rafet Sifa, Christian Bauckhage, Hannaneh Hajishirzi, and Yejin Choi. Is reinforcement learning (not) for natural language processing: Benchmarks, baselines, and building blocks for natural language policy optimization, 2023.

[30] Chamani Shiranthika, Kuo-Wei Chen, Chung-Yih Wang, Chan-Yun Yang, BH Sudantha, and Wei-Fu Li. Supervised optimal chemotherapy regimen based on offline reinforcement learning. IEEE Journal of Biomedical and Health Informatics, 26(9):4763–4772, 2022.

[31] Laura Smith, Ilya Kostrikov, and Sergey Levine. A Walk in the Park: Learning to Walk in 20 Minutes With Model-Free Reinforcement Learning, August 2022.

[32] Charlie Snell, Ilya Kostrikov, Yi Su, Mengjiao Yang, and Sergey Levine. Offline rl for natural language generation with implicit language q learning. arXiv preprint arXiv:2206.11871, 2022.

[33] Matthijs TJ Spaan. Partially observable markov decision processes. In Reinforcement learning: State-of-the-art, pages 387–414. Springer, 2012.

[34] Denis Tarasov, Alexander Nikulin, Dmitry Akimov, Vladislav Kurenkov, and Sergey Kolesnikov. Corl: Research-oriented deep offline reinforcement learning library. arXiv preprint arXiv:2210.07105, 2022.

[35] Amol Thakkar, Veronika Chadimová, Esben Jannik Bjerrum, Ola Engkvist, and Jean-Louis Reymond. Retrosynthetic accessibility score (rascore)–rapid machine learned synthesizability classification from ai driven retrosynthetic planning. Chemical Science, 12(9):3339–3349, 2021.

[36] Oleg Trott and Arthur J Olson. Autodock vina: improving the speed and accuracy of docking with a new scoring function, efficient optimization, and multithreading. Journal of computational chemistry, 31(2):455–461, 2010.

[37] Ashish Vaswani, Noam Shazeer, Niki Parmar, Jakob Uszkoreit, Llion Jones, Aidan N Gomez, Łukasz Kaiser, and Illia Polosukhin. Attention is all you need. Advances in neural information processing systems, 30, 2017.

[38] Izhar Wallach, Michael Dzamba, and Abraham Heifets. Atomnet: A deep convolutional neural network for bioactivity prediction in structure-based drug discovery. CoRR, abs/1510.02855, 2015. URL http://arxiv.org/abs/1510.02855.

[39] Chenran Wang, Yang Chen, Yuan Zhang, Keqiao Li, Menghan Lin, Feng Pan, Wei Wu, and Jinfeng Zhang. A reinforcement learning approach for protein–ligand binding pose prediction. BMC bioinformatics, 23(1):1–18, 2022.

[40] David Weininger. Smiles, a chemical language and information system. 1. introduction to methodology and encoding rules. Journal of Chemical Information and Computer Sciences, 28(1):31–36, 1988. doi: 10.1021/ci00057a005. URL 10.1021/ci00057a005.

[41] Shenghao Wu, Tianyi Liu, Zhirui Wang, Wen Yan, and Yingxiang Yang. Rlcg: When reinforcement learning meets coarse graining. In NeurIPS 2022 AI for Science: Progress and Promises, 2022.

[42] Jiaxuan You, Bowen Liu, Zhitao Ying, Vijay Pande, and Jure Leskovec. Graph convolutional policy network for goal-directed molecular graph generation. Advances in neural information processing systems, 31, 2018.

[43] Tengteng Zhang and Hongwei Mo. Reinforcement learning for robot research: A comprehensive review and open issues. International Journal of Advanced Robotic Systems, 18(3):17298814211007305, 2021. doi: 10.1177/17298814211007305. URL 10.1177/17298814211007305.

